# What makes *Hemidactylus* invasions successful? A case study on the island of Curaçao

**DOI:** 10.1101/2020.04.17.047209

**Authors:** April D. Lamb, Catherine A. Lippi, Gregory J. Watkins-Colwell, Andrew Jones, Dan Warren, Teresa L. Iglesias, Matt Brandley, Connor Neagle, Alex Dornburg

**Affiliations:** North Carolina Museum of Natural Sciences, Raleigh, NC 27601; North Carolina State University, Department of Applied Ecology, Raleigh, NC 27695; Quantitative Disease Ecology and Conservation (QDEC) Lab Group, Department of Geography, University of Florida, Gainesville, FL 32611; Division of Vertebrate Zoology, Yale Peabody Museum of Natural History, New Haven, CT 06520 USA; Department of Ecology and Evolutionary Biology, Yale University, New Haven, CT 06520 USA; Senckenberg Biodiversität und Klima - Forschungszentrum, Frankfurt am Main 60325, Germany; Okinawa Institute of Science and Technology Graduate University, Okinawa Prefecture 904-0495, Japan; School of Biological Sciences, University of Sydney, NSW 2006, Australia; North Carolina State University, Department of Forestry and Environmental Resources, Raleigh, NC 27695

**Keywords:** invasive species, urbanization, vertebrate biodiversity loss, food web, trophic ecology

## Abstract

*Hemidactylus* spp. (House geckos) rank among the most successful invasive reptile species worldwide. *Hemidactylus mabouia* in particular has become ubiquitous across tropical urban settings in the Western Hemisphere. *H. mabouia’s* ability to thrive in close proximity to humans has led to the rapid displacement of native geckos in urban areas, however the mechanisms driving this displacement remain understudied. Here we combine data from nitrogen and carbon stable isotopes, stomach contents, and morphometric analyses of traits associated with feeding and locomotion to test alternate hypotheses of displacement between *H. mabouia* and a native gecko, *Phyllodactylus martini*, on the island of Curaçao. Consistent with expectations of direct food resource competition, we demonstrate substantial overlap of invertebrate prey resources between the species. Additionally, we found strong evidence from both diet content and stable isotope analyses that *H. mabouia* acts as a vertebrate predator, preying upon *P. martini* as well as other native and non-native reptiles. Finally, we show that *H. mabouia* possesses several morphological advantages, including larger sizes in feeding-associated traits and limb proportions that could offer a propulsive locomotor advantage on vertical surfaces. Together, these findings suggest the successful establishment of *H. mabouia* likely involves a combination of both exploitative interspecific competition and predation. Given the ubiquity of *H. mabouia*, illuminating the role of this species as both a competitor and a predator casts new concerns on the ecological and demographic impacts of this widespread urban invader.

## Introduction

Since the onset of the industrial revolution, the impact of invasive species on endemic fauna and flora has been a central topic in the management and conservation of biodiversity worldwide (Paini et al. 2016; Young et al. 2017; Shechonge et al. 2019). This concern reflects dramatic losses in global biodiversity and an increasing shift towards widespread homogenization of the planet’s biota (McKinney and Lockwood 1999; McKinney 2006; Trentanovi et al. 2013). These trends are especially acute in urbanizing landscapes, which have repeatedly been shown to support higher numbers of non-native, human-commensal species (Useni Sikuzani et al. 2018), such as cats (Buzan 2017; Bateman and Fleming 2012), rats (Bateman and Fleming 2012; Buzan 2017), and house sparrows (González-Oreja et al. 2018). Following establishment, successful non-native species have been found to restructure resident community assemblages by directly or indirectly altering top-down processes (e.g. predation, (Willson 2017; Pedersen et al. 2018)), bottom-up processes (e.g. resource availability (Yam et al. 2016)), or both (i.e., “middle-out” effects, (Weber and Brown 2009)) at the expense of native taxa. In the most extreme cases this can result in the extirpation or extinction of native species (Wiles et al. 2003; Toussaint et al. 2016; Liu et al. 2017). However, investigations into the impact and distribution of introduced species have been largely restricted to species that are easily visible in the landscape (Beasley et al. 2018), are a direct nuisance to humans (Bithas et al. 2018), or displace commercially important or game species (Galanidi et al. 2018; Hill et al. 2004). While not misguided, this bias has left a critical gap in our knowledge regarding the potential impacts of less readily observable, but equally common, non-native human-commensal taxa (Morais and Reichard 2018).

Despite the prevalence of invasive reptiles around the world (Kraus 2015), most attention has been devoted to the loss of biodiversity following the spread of a few larger bodied species such as Burmese pythons (Smith et al. 2016; Willson 2017), green iguanas (Falcón et al. 2013; Burgos-Rodríguez et al. 2016), and brown tree snakes (Wiles et al. 2003; Rodda and Savidge 2007; Richmond et al. 2015). However, numerous smaller and more clandestine reptiles have also become globally pervasive (Kraus 2015; Capinha et al. 2017; Lapiedra et al. 2017). These invasions, while common, often go unnoticed until native reptiles begin to disappear from the landscape (Kraus 2015). Such cryptic losses in biodiversity are a hallmark of introduction of *Hemidactylus* spp. (House Geckos), a group commonly associated with urbanized and developing areas. Over the past century, *Hemidactylus* spp. have become an established feature of tropical and subtropical landscapes around the world (Carranza and Arnold 2006). Following establishment, these geckos have been repeatedly linked to local extirpation and even extinction of native lizards (Petren and Case 1996; Cole et al. 2005; Hoskin 2011). One species in particular, *Hemidactylus mabouia* (Tropical House Gecko), is perhaps the most pervasive and formidable gecko to invade the Western Hemisphere (Weterings and Vetter 2018).

Native to Africa, *Hemidactylus mabouia* is now common throughout the Americas and Caribbean (Carranza and Arnold 2006). Recent studies have linked the successful establishment of this species in urban and suburban environments to its ability to capitalize on the aggregation of insects around human light sources (Hughes et al. 2015). Restricting the access of native geckos to these clustered food resources is thought to represent a competitive advantage for *H. mabouia* that promotes high densities of individuals (van Buurt 2004; Short and Petren 2011; Williams et al. 2016). As *H. mabouia* adult males are noted for being particularly aggressive (Short and Petren 2011), the ability of this species to aggressively restrict access to spatially clustered food resources suggests interference competition, whereby high densities of aggressive competitors fuel the displacement of native gecko species. However, alternate hypotheses remain untested.

In addition to their impact as competitors, two aspects of *H. mabouia* invasions that have received particularly little attention are locomotor morphology and role of *H. mabouia* as potential predators. The feeding mode of *H. mabouia* combines ambush tactics (Vitt 1983) with active pursuit of nearby prey (Dornburg et al. 2016). Such a foraging mode could have selected for limb proportions that offer a mechanical advantage on sheer vertical surfaces (Zaaf and Van Damme 2001). Further, it is possible that the generally robust body plan of *Hemidactylus* spp. facilitates the capture of larger prey not accessible to other similarly sized geckos, although this hypothesis has not been tested. An alternative, but not mutually exclusive, explanation for the success of *H. mabouia* comes from isolated natural history reports of *H. mabouia* preying on other species of geckos (Dornburg et al. 2011, 2016) as well as cannibalizing conspecifics (Bonfiglio et al. 2006). Both morphological advantages and predation have been invoked as major drivers of displacement in the wake of invasions by the closely related *Hemidactylus frenatus* in the Pacific (Petren and Case 1998; Petren and Case 1996; Bolger and Case 1992; Case et al. 1994; Short and Petren 2012). However, both the role of morphological advantages and predation in driving the decline of native gecko populations in the wake of an *H. mabouia* invasion remain unclear.

In this study we assess whether there is evidence for trait driven advantages or predation in the invasion of *H. mabouia* in the Lesser Antillean island of Curaçao. We specifically focus on competition between *H. mabouia* and the native *Phyllodactylus martini* (Dutch Leaf Tailed Gecko), as *P. martini* declines have historically been linked to the invasion of *H. mabouia* (van Buurt 2004; van Buurt 2006; Hughes et al. 2015; Dornburg et al. 2016). First we integrate analyses of nitrogen and carbon stable isotopes with direct examination of stomach contents to test for levels of prey overlap and isotopic trophic signatures consistent with hypotheses of resource competition or predation. We additionally collected morphometric measurements from traits associated with feeding and locomotion to test the hypothesis that *H. mabouia* possess trait advantages over its hypothesized native competitor.

## Materials/Methods

### Fieldwork and Data Acquisition

*Hemidactylus mabouia* (n=90) and *Phyllodactylus martini* (n=71) specimens were collected at six sites across Curaçao between July 2009 and September 2011: Lagun, Wespunt, CARMABI, Shete Boca, Saint Anna Bay, and Willemstadt (Figure 1; Supplemental materials). Habitat type and species occupancy vary across sampling locations. For example, both species co-occur in Lagun and Westpunt. At these sites we restricted our sampling to suburban areas near natural habitats to maximize the potential of both species co-occurring as *P. martini* has been found to be absent far from edge habitats in the presence of *H. mabouia* (Hughes et al. 2015). In contrast, Shete Boca is a natural area in which *H. mabouia* are absent, while Saint Anna Bay and Willemstadt are urban areas in which *P. martini* are absent. This sampling design allowed us to capture a greater degree of diet breadth of each species across the island. Across sites, sample locations included walls, rocks, outcrops, trees, thatch roofs, open ground, and shrubbery. At no point during sampling did we document individuals of both species occupying the same structure (e.g., same wall or tree), and individuals were collected opportunistically at each site. Prior to preservation, muscle biopsies were taken from each individual and dehydrated for analysis of stable isotopes. Additionally, leaf samples from each locality and temporal sampling event were collected and dehydrated for use as baselines in isotopic analyses. Specimens then were fixed in 10% formalin and later transferred to 70% ethanol and deposited in the Yale Peabody Museum of Natural History (supplemental materials).

**Fig. 1.**
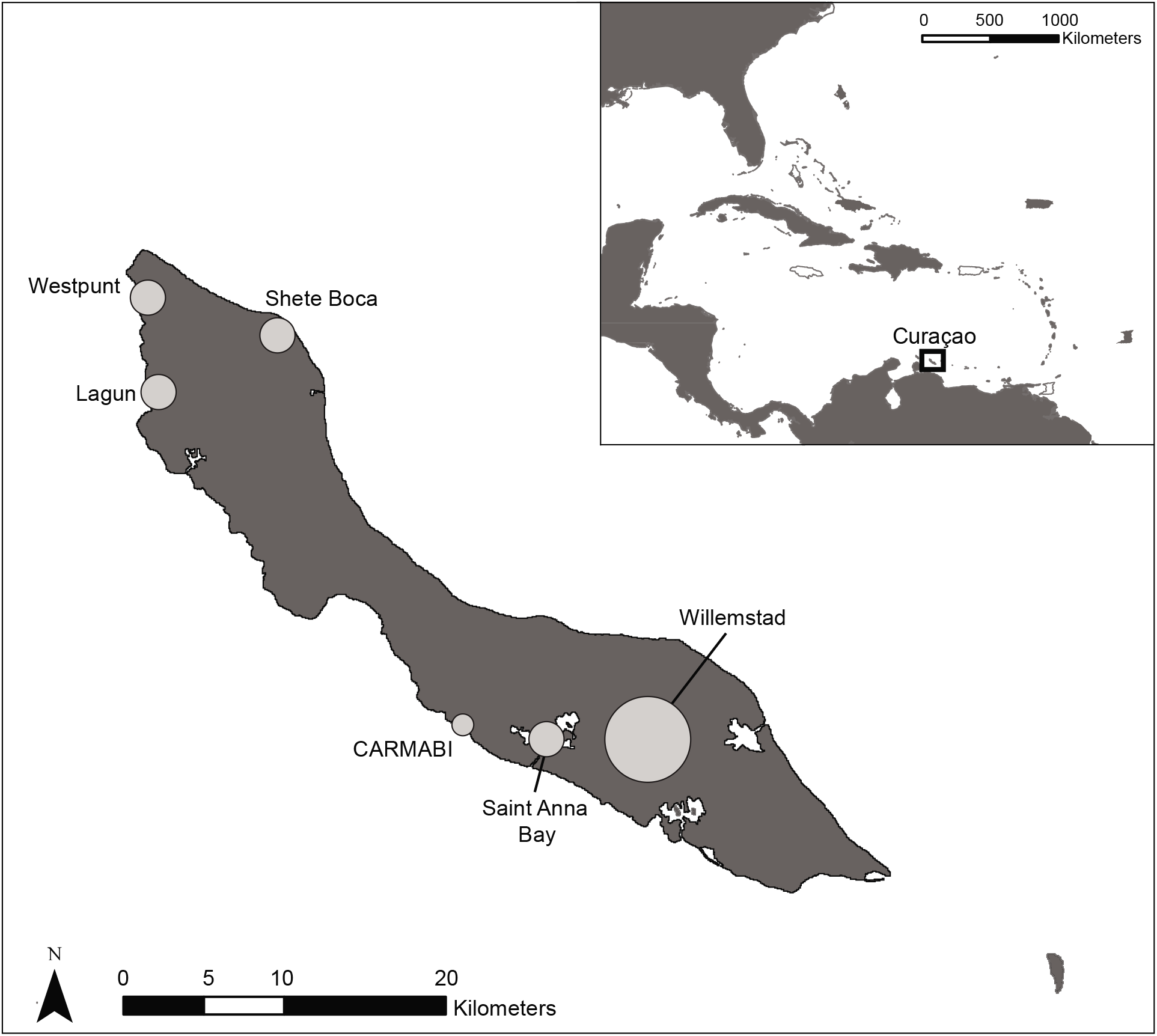
Map of study area. Circles indicate sampling locations.

Specimens collected in 2011 (n=59) had their stomach contents preserved in 10% formalin and dissected, with contents identified and enumerated under a dissecting MVX10 microscope (Olympus Corp.; http://www.olympus-lifescience.com/). Prey items were identified to the taxonomic groupings similar to those in other studies of Caribbean lizards (e.g., (Perry 1996)): Arachnida (scorpiones), Arachnida (Araneae), Blattaria (Blattodea), Chilopoda, Coleoptera, Diptera, Ephemeroptera, Hemiptera, Hymenoptera, Isopoda, Lepidoptera, Orthoptera, and “other”. Any vertebrate remains encountered were additionally identified to the highest taxonomic resolution possible, and we additionally identified any parasites encountered in the stomach. As formalin and alcohol preservation can have heterogeneous effects on the volume of invertebrate organisms (Donald and Paterson 1977), enumeration of diet contents was restricted to % frequency.

We further collected measurement data on 79 *Hemidactylus mabouia* for 10 morphological traits associated with feeding and locomotion: snout-vent length (SVL), postorbital width, temporalis width, head length, jaw length, head height, humerus length, radius length, femur length and tibia length. All measurements were taken to the nearest 0.01 mm using digital calipers (Fowler Promax). Both stomach content and morphological data were integrated with the dataset of Dornburg et al. (2016) who previously measured *Phyllodactylus martini* specimens for the same morphological traits (n=34); Zenodo DOI: 10.5281/zenodo.61569) and prey items (n=69; Zenodo DOI: 10.5281/zenodo.61569).

### Stable isotopic analysis of trophic ecology

Leg muscle biopsies from 21 individual *Hemidactylus mabouia* and 17 *Phyllodactylus martini* legs as well as 8 plant stems and leaf baseline samples were used in nitrogen (∂15N) and carbon (∂13C) stable isotope analysis. Skin was removed from each muscle biopsy, and individual muscle and plant baseline samples were dehydrated at 40°C degrees for 48 hours. Following dehydration, samples were powdered using a bead beater (MP FastPrep24 Hyland Scientific). From each sample, 1.5 mg of powder was loaded into 3×5 mm tins. Samples were analyzed at the University of California Davis Stable Isotope Facility using an isotope ratio mass spectrometer. As nitrogen enrichment can vary over spatial or temporal periods, quantification of trophic position for each individual was standardized using primary producer baseline samples from plant leaves and stems collected at each locality (Vidal and Sabat 2010; Roches et al. 2016). To account for ∂15N values not reflecting primary producer level values (Marshall et al. 2007), baseline samples were compared across sites with aberrant samples (i.e., primary producer ∂15N > consumer ∂15N) removed. Nitrogen values were standardized following Post (2002), in subtracting the mean ∂15N of the primary producers from ∂15N of each individual lizard and assuming fractionation of 3.4% per trophic level (Post 2002). ∂15N values for each species were visualized using violin plots which allow for simultaneous inspection of quartiles and the underlying probability distribution through integration of a rotated kernel density plot with a boxplot (Hintze and Nelson 1998). We tested for differences between the mean ∂15N values of *H. mabouia* and *P. martini* using a Welch’s t-test and additionally used Levene’s test to assess whether there was a significant difference in ∂15N variance between species. A significant positive difference in ∂15N between *H. mabouia* and *P. martini* would be consistent with the hypothesis that individual *H. mabouia* are vertebrate predators. To test for potential differences in ∂13C, we used the same statistical approaches as those used in the analysis of ∂15N, assuming carbon fractionation to be 0% (Post 2002). In this case, non-significant differences in ∂13C would support the expectation that these species forage in similar habitats. All analyses were conducted in R, v. 3.4.3 (R Development Core Team 2018).

### Stomach Content Analysis

For each species, relative frequency of each prey item was calculated based on the total number of prey items encountered across all individuals of that species. Differences in stomach contents between species were visualized using a non-metric multidimensional scaling (NMDS) analysis in the vegan software package (Oksanen et al. 2007; Oksanen 2011). Stress values were quantified to test if NMDS ordination represents a viable indicator of species dissimilarity, with stress values less than 0.1 indicating good representation of the dissimilarities (Clarke 1993). An analysis of similarity (ANOSIM; (Clarke 1993); (Chapman and Underwood 1999)) was used to additionally test for significant differences between species, using Manhattan distances and 999 permutations in the vegan software package (Oksanen et al. 2007; Oksanen 2011). Differences in mean ranks were quantified using the R statistic, with values close to zero indicating high similarity and values close to one indicating high dissimilarity (Chapman and Underwood 1999). As *P. martini* has been found to consume different prey in rural versus suburban edge habitats (Dornburg et al. 2016), NMDS and ANOSIM analyses were repeated with analyses restricted to individual geckos collected in areas where both species co-occur. This allowed us to test whether pooling across habitat types potentially masked differences or overlap in prey items. Additionally, differences in parasite load between species were compared using a Welch’s t-test.

### Comparisons of morphology

We compared absolute differences in log snout-vent length (SVL) between species using an ANOVA and created raincloud plots (Allen et al. 2019) to visualize differences. These plots combine classic boxplots with violin raw data plots to simultaneously visualize data, the difference in size quartiles, and a kernel density estimate of the probability density of the SVL data. We conducted a principal components analysis (PCA) to visualize the overall morphospace occupied by both species. In geckos, size has been shown to covary with our target morphological measurements (Dornburg et al. 2016). As such, we first regressed all the measurements per species against SVL (supplemental materials) and used the residual values of individual traits regressed against SVL as data for the PCA. To assess if differences in morphospace occupancy were mostly driven by uneven sample sizes, we randomly sampled equal numbers of both gecko species from our data 200 times in intervals of 5 between 10 and 55. For each of these 2000 datasets, we conducted a PCA and computed the mean and quantiles (25% & 75%) of the ratio of *H. mabouia* to *P. martini* morphospace.

While morphospace visualization is advantageous for assessing the overall overlap of phenotypic variation, it is possible that allometric slopes are identical between species and simply have different intercepts (i.e., at a given body size a focal trait in one species is larger in one species than the other). To further scrutinize our data, we used an analysis of covariance (ANCOVA) to test for differences in each morphological trait between species. For each analysis, we kept log transformed SVL as the covariate and treated each log transformed morphometric measurement (e.g., jaw length, limb length, etc) as the response. This approach allowed us to test the potential correlation for each measured trait and SVL as well as the possibility of significant differences between species that take trait covariation with SVL into account. We repeated analyses with non-significant interactions removed, as inclusion or omission of non-significant interactions can potentially impact ANCOVA analyses.

In many lizard species, including geckos, head size is a sexually dimorphic trait with males often having larger heads relative to females (Kratochvíl et al. 2002; Scharf and Mieri 2013; Iturriaga and Marrero 2013). As such, we used an ANCOVA to assess whether morphological differences for each trait were potentially masked when pooling sexes by species. For all analyses, we again kept log transformed SVL as the covariate and treated each log transformed morphometric measurement as the response. Finally, we assessed potential differences in total limb lengths (humerus length + radius length; femur length + tibia length) between species and sexes using log transformed limb length as the response and log transformed SVL as the covariate in an ANCOVA. This additional analysis facilitated additional comparisons of expectations of gecko locomotion as studies often discuss differences in total limb lengths.

Prior work has suggested large hind limbs compensate for large heads in the locomotion of *Hemidactylus* spp. geckos (Cameron et al. 2013). As such, we examined scaling relationships between head size and hindlimb length for both species by constructing a set of generalized linear models (GLMs). We built models using sex, species, SVL, and head size as explanatory variables, with one set of models using head length to quantify head size and another set using post-orbital width. All models except the intercept-only null models contained an interaction term between SVL and the head size term, so that the effects of head size on limb length would be controlled for overall body size. Additional candidate models included (1) sex, (2) species identity, and (3) sex, species identity, and an interaction term between species identity and head size. Model fit was evaluated with the Akaike information criterion with a correction for small sample size (AICc). This method of model selection identifies models that predict the data well while penalizing overparameterization (Burnham and Anderson 2004).

## Results

### Differences in feeding ecology

Analysis of ∂15N revealed a significant (Welch’s t-test: p < 0.004; t= 3.123 df = 34.272) shift towards a higher mean trophic level in *H. mabouia* versus *P. martini* (Figure 2A). In contrast, analysis of ∂13C isotopes revealed no significant (Welch’s t-test: p < 0.401; t= 0.857 df = 21.812) shift in mean ∂13C between *H. mabouia* versus *P. martini* (Figure 2B), supporting the expectation that these two species overlap in major foraging habitat type. Levene’s tests did not support a significant increase in ∂15N variance within *H. mabouia* versus *P. martini* (F= 0.480; p = 0.493), though a single *P. martini* outlier point in our analysis depicted a carbon signature consistent with marine prey resource use, suggesting the possibility that some individuals may opportunistically forage close to the shoreline. Regardless, the difference in variance of ∂13C values between species was found to be non-significant (F= 0.585; p=0.449), even after removing this potential outlier point (F= 7.624; p=0.155).

**Fig. 2.**
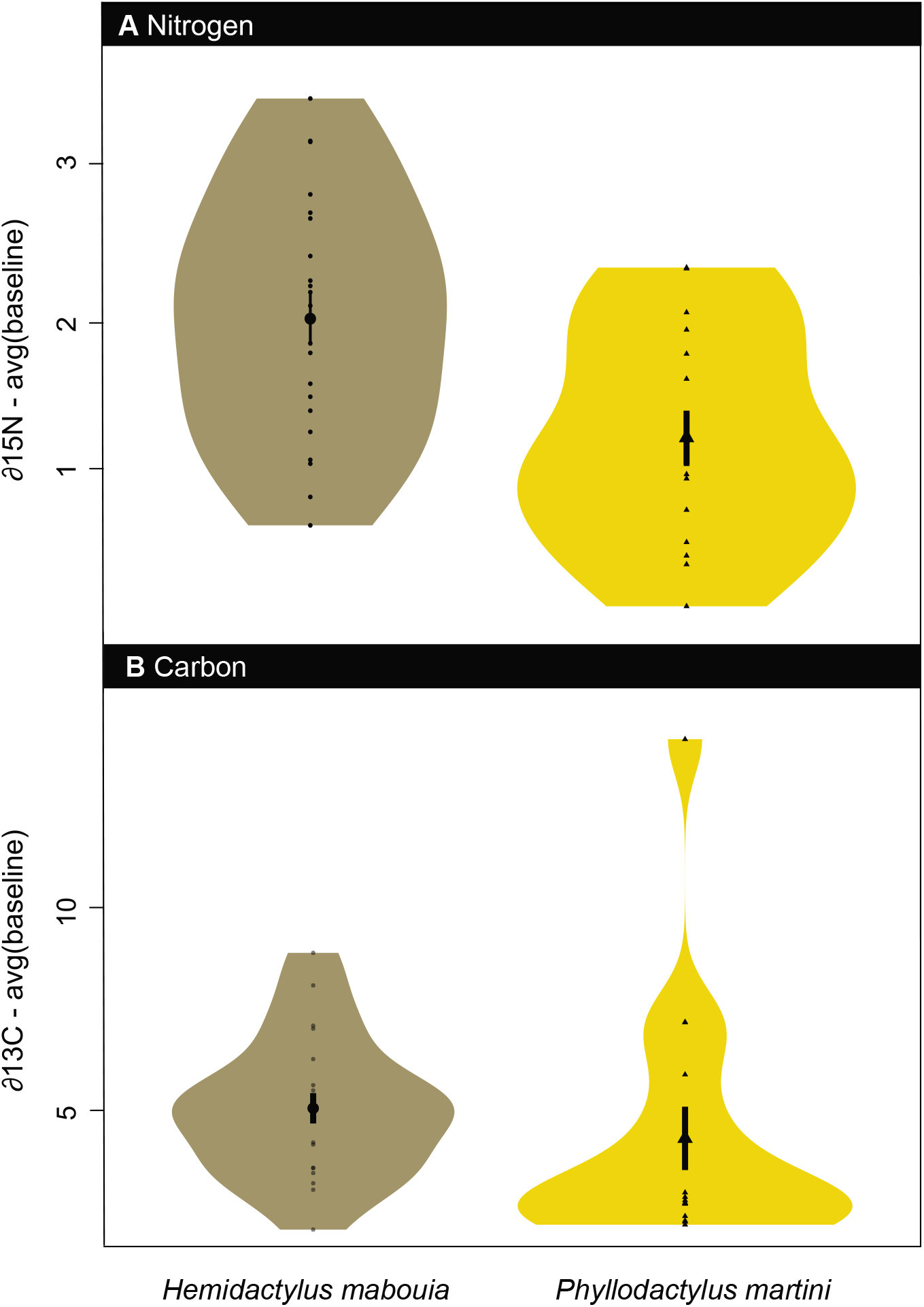
Violin plots of isotopic data. **A.** Estimated trophic position for *Phyllodactylus martini* (n=17) and *Hemidactylus mabouia* (n=21) using Nitrogen. **B.** Carbon. Raw nitrogen and carbon isotopic values were corrected using average baseline values across all sites.

Our analyses of individual stomach contents revealed *H. mabouia* to generally have fewer prey items per stomach than *P. martini* (Welch’s t-test: p < 0.001; t= 3.31 df = 84.74). Across 59 specimens of *H. mabouia*, we found 0 to 3 prey items per individual, which spanned a wide range of invertebrates (Table 1 & Figure 3). Additionally, three individual *H. mabouia* each contained a single vertebrate prey item. These prey items were identified as *Gonatodes antillensis*, *Phyllodactylus martini*, and *Ramphotyphlops braminus*. Comparing the invertebrate prey found in *H. mabouia* to *P. martini* revealed the two species to consume similar prey items with differences in the overall percentages of prey items consumed. We find both species to generally consume the same major invertebrate prey groups but at different frequencies: Arachnida (*H. mabouia* = 14%; *P. martini* =19%); Insecta (*H. mabouia* = 50%; *P. martini* =58%); Isopoda (*H. mabouia* = 28%; *P. martini* =9%). Further, the species varied with regard to individual prey items and frequency within these major groupings (Figure 3 & Table1) and ANOSIM results supported significant differences between groups (R=0.213, p=0.001). Visualizations of diet data based on NMDS analyses of invertebrate prey items for the two species support a large degree of overlap in diet with *H. mabouia* utilizing the same resources as *P. martini*, but with *H. mabouia* also utilizing more resources not exploited by *P. martini* (Figure 4A). Repeating analyses for just individuals residing in areas of co-occurrence again supported significant differences between species in an ANOSIM analysis when vertebrates were included as a prey category (R=0.020, p=0.040). All instances of vertebrate predation by *H. mabouia* were found in areas where the two species overlap (supplemental materials). Restricting an ANOSIM analyses to just invertebrate prey items supported no significant differences in diet between the two species (R=0.018, p=0.070). Visualizations of both the raw (supplemental materials) and NMDS analyses of the invertebrate prey item diet data for the two species areas further depicted a large degree of overlap (Figure 4B). In addition to prey contents, parasitism infestations by nematodes were significantly different between the two species (Welch’s t-test: p < 0.001; t= −3.768 df = 71), suggesting higher parasite pressure within *P. martini* (Table 1).

**Fig. 3.**
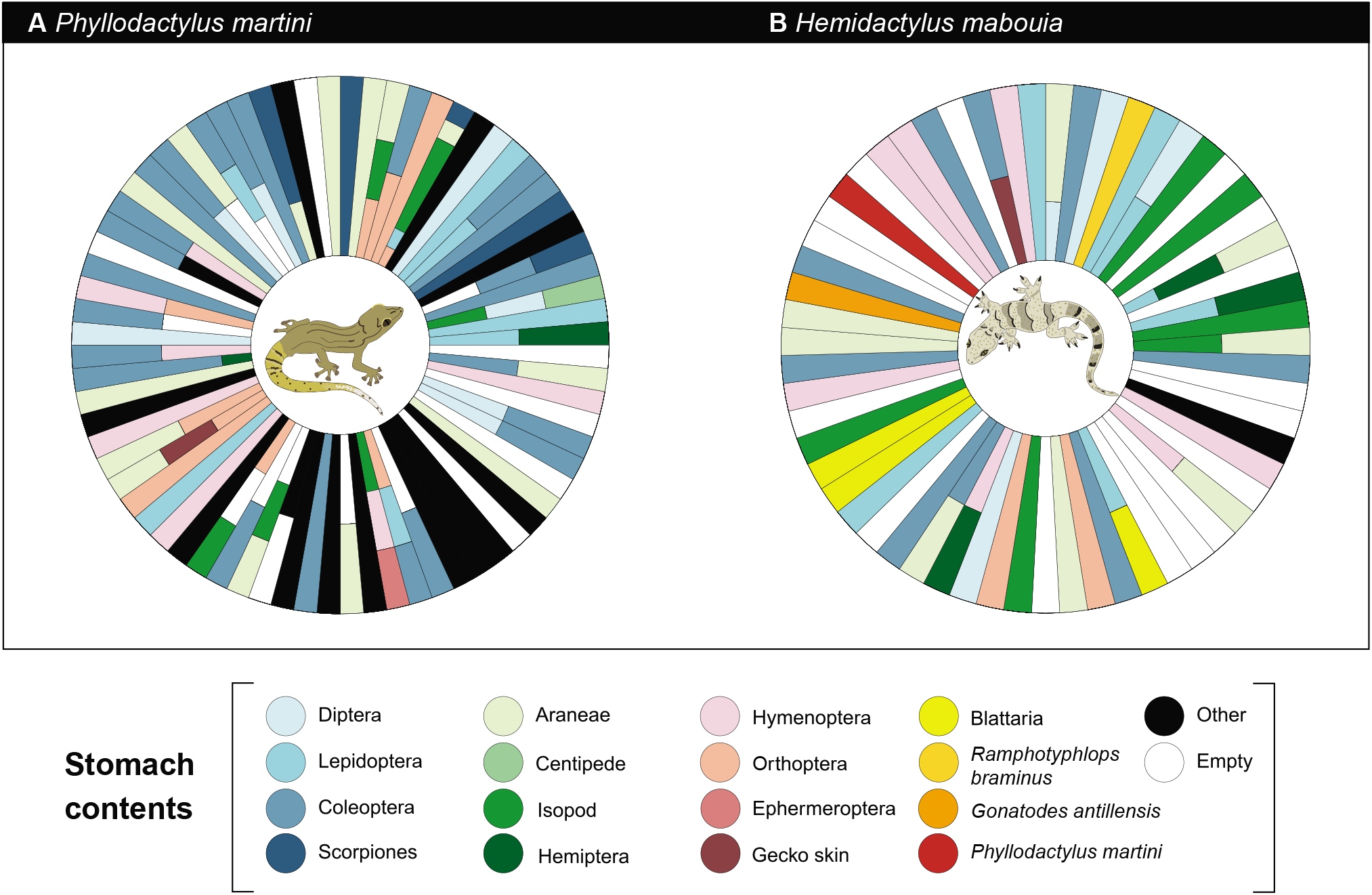
Visualization of stomach content data. **A.** Data from analyzed *Phyllodactylus martini* (n=72). **B.** *Hemidactylus mabouia* (n=59). Columns in sphere correspond to the relative frequency of an individual’s prey items. Colors correspond to matching prey categories in legend.

**Fig. 4.**
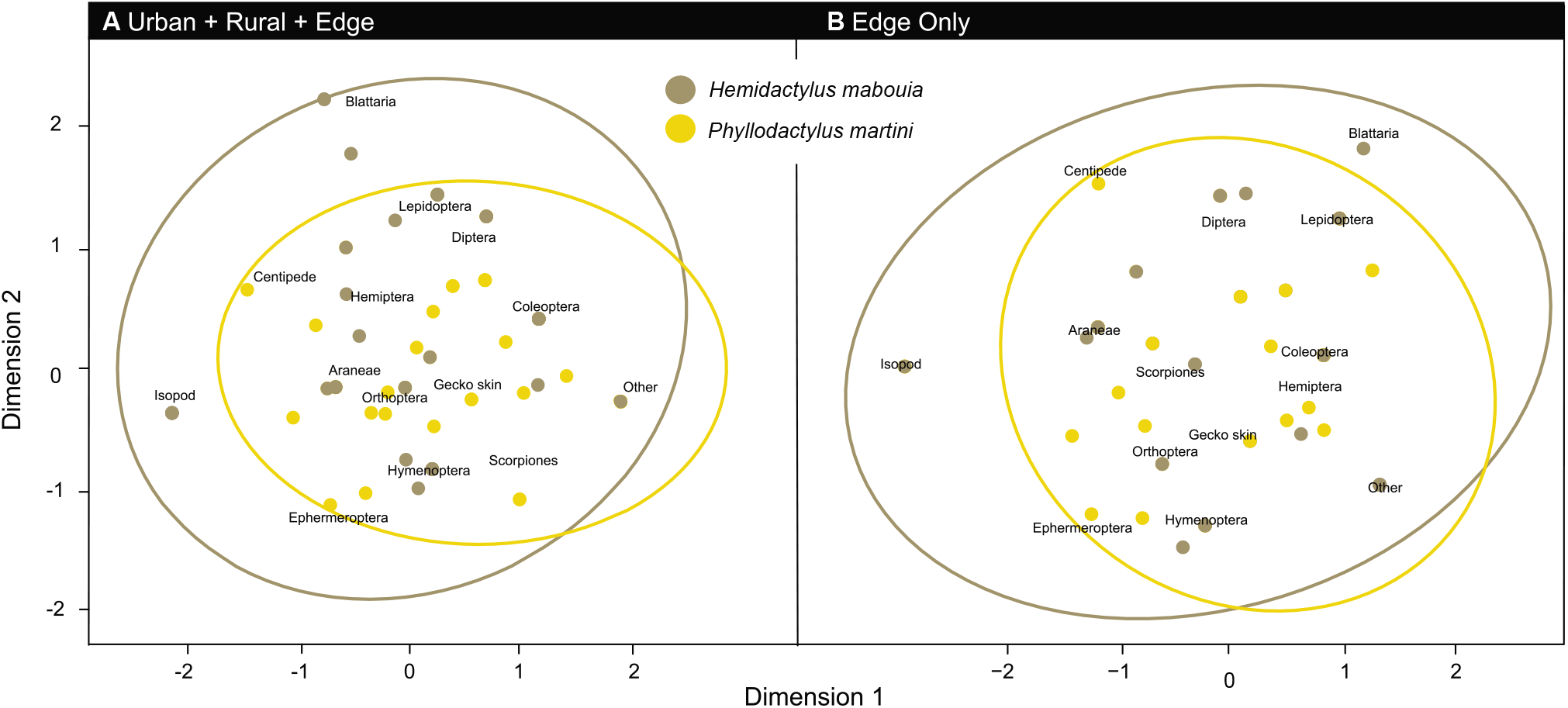
Non-metric multidimensional scaling analysis contrasting stomach contents. Data from analyzed *Phyllodactylus martini* (n=79) and *Hemidactylus mabouia* (n=57). Ellipses encompass the 95% confidence interval around the centroid of each species, and prey labels indicate location of prey categories within the diet space.

**Table 1.** Relative frequency of prey stomach content items across all sampled individual *Hemidactylus mabouia* (total items = 72) and *Phyllodactylus martini* (total items = 115). Bold values indicate sums of prey items within major categories (i.e., Insecta).

|  | *Hemidactylus mabouia* | *Phyllodactylus martini* |
| --- | --- | --- |
| Prey | Relative Frequency | Relative Frequency |
| **Arachnida** | **0.141** | **0.191** |
| Scorpiones | --- | 0.052 |
| Araneae | 0.14 | 0.139 |
| **Chilopoda** | --- | **0.009** |
| **Insecta** | **0.507** | **0.583** |
| Blattaria | 0.042 | 0.00 |
| Coleoptera | 0.127 | 0.235 |
| Diptera | 0.056 | 0.078 |
| Ephemeroptera | 0 | 0.009 |
| Hemiptera | 0.056 | 0.017 |
| Hymenoptera | 0.099 | 0.061 |
| Lepidoptera | 0.099 | 0.096 |
| Orthoptera | 0.028 | 0.087 |
| **Isopoda** | **0.282** | **0.087** |
| **Skin Shed** | **0.014** | **0.009** |
| **Vertebrata** | **0.042** | --- |
| *Gonatodes antillensis* | **0.014** | --- |
| *Phyllodactylus martini* | **0.014** | --- |
| *Ramphotyphlops braminus* | **0.014** | --- |
| **Other** | **0.014** | **0.128** |

### Differences in morphology

We found a significant overall size difference between *H. mabouia* versus *P. martini* (F= 10.61; p = 0.00143), with *H. mabouia* generally being larger (Figure 5A). Three axes of a principal components analysis (PCA) of morphological traits collectively capture 64.1% of the measured variation (PC1: 34.84%; PC2: 16.90%; PC3: 12.37%). PC1 largely captures differences in limb lengths (~39% total hindlimb, 18% total front limb) and variation in the postorbital width (~24%). In contrast, PC2 mostly captures variation in cranial measurements with over 70% of the loadings belonging to a combination of head length (~29%), jaw length (~17%), temporalis width (~13%), and postorbital width (~13%). PC3 largely captured further variation in cranial morphology (Supplemental materials). Visualization of these PC axes revealed a high degree of overlap between species, with *H. mabouia* occupying more morphospace overall. Between PC1 and PC2 (Figure 5B) the total morphospace occupancy based on the convex hull area [CHA] of *H. mabouia* was 64% larger (*H. mabouia* CHA = 18.370; *P. martini* = 11.140). Similarly, between PC1 & PC3 (*H. mabouia* CHA = 22.724; *P. martini* = 5.898; Figure 5C) and PC2 & PC3 (*H. mabouia* CHA = 16.532; *P. martini* = 5.474; Figure 5D) the CHAs of *H. mabouia* were larger. Results of our dataset resampling analyses support that these differences were not due to sample size differences alone (Supplemental materials). SVL was significantly correlated with all measured morphological traits (Table 2; Supplemental materials) and ANCOVA results further support significant differences between residual trait variation after accounting for SVL scaling between species for all traits (Table 2; Supplemental materials). The only exception to this general trend of a significant relationship between species identity and trait was head height (F= 3.232; p= 0.075). These results were consistent whether non-significant interactions were included in the analysis or not (Supplemental materials). Tests for sexual dimorphism for no evidence for trait differences between male and female *P. martini*. In contrast, head width was significantly different between male and female *H. mabouia*, suggesting *H. mabouia* males have wider heads than females (Supplemental materials).

**Fig. 5.**
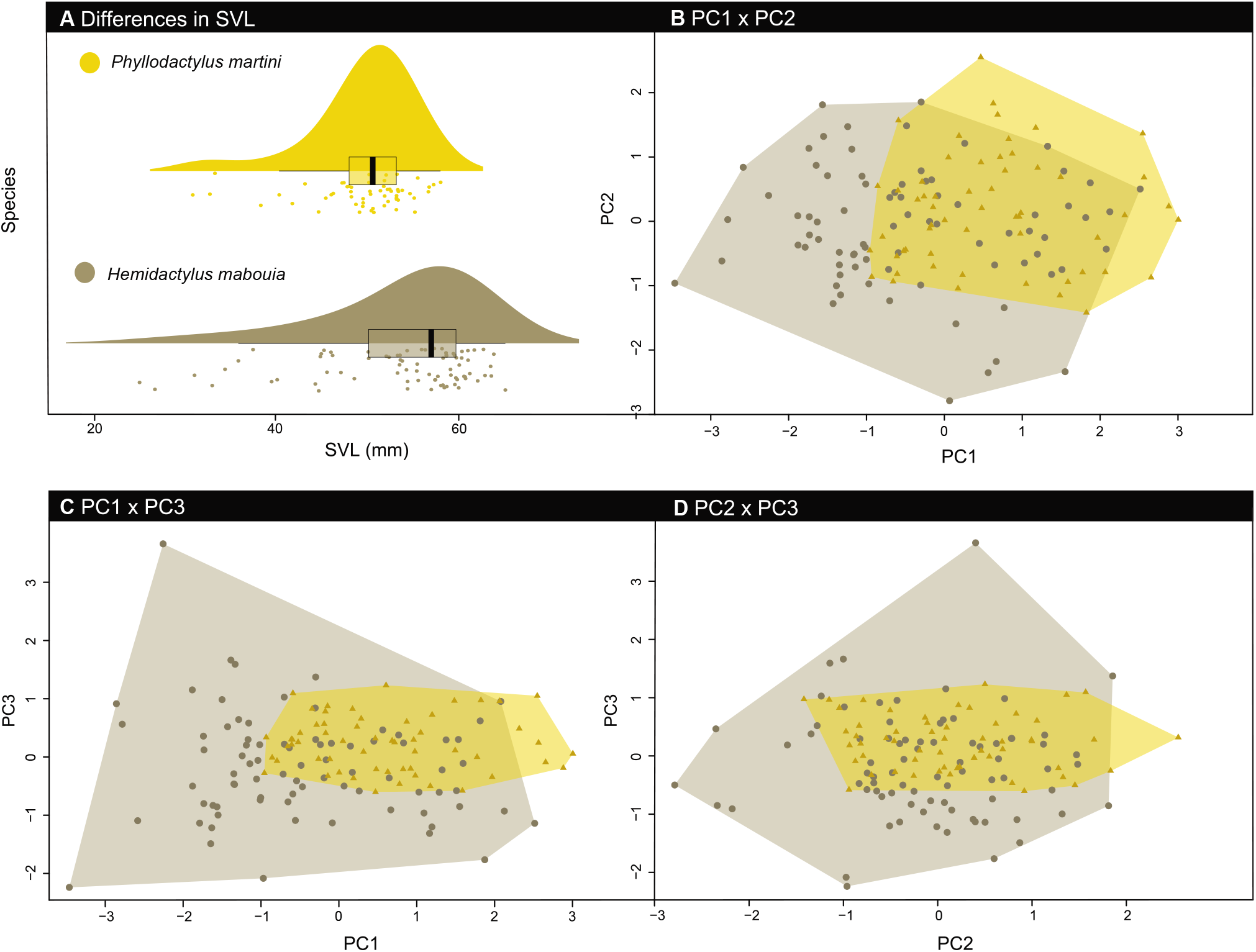
Analysis of morphometric traits. **A.** Raincloud plots visualizing SVL differences between *Phyllodactylus martini* (yellow) and *Hemidactylus mabouia* (brown), depicting the probability distribution through a rotated violin plot (top), box plot summary of quartiles (middle), and raw data (bottom) for each species. **B-D.** Principal components analysis showing overlap of morphological traits between species. Principal component scores are visualized for each axis and species with background shading representing fitted convex hulls of the morphospace occupied by each species.

**Table 2.** ANCOVA results testing the effect of snout-vent length (SVL), species, and their interaction on measured morphological characters. Bolded values indicate significant effects. * stands for *P*-values ranging from 0.05 to 0.01, ** for *P*-values ranging from 0.01 to 0.001 and *** for *P*-values smaller than 0.001

| Trait | log(SVL) (F/P) | Species (F/P) | Species:log(SVL) (F/P) |
| --- | --- | --- | --- |
| Post Orbit Width | **121.82/\*\*\*** | **5.98/\*** | 0.07/0.78 |
| Temporalis Width | **1151.42/\*\*\*** | **32.45/\*\*\*** | 0.07/0.78 |
| Head Length | **1298.73/\*\*\*** | **101.84/\*\*\*** | 0.07/0.78 |
| Jaw Length | **526.77/\*\*\*** | **86.16/\*\*\*** | 0.18/0.66 |
| Head Height | **358.09/\*\*\*** | 3.23/0.07 | 0.35/0.55 |
| Humerus Length | **163.53/\*\*\*** | **43.52/\*\*\*** | **8.30/\*\*** |
| Radius Length | **464.95/\*\*\*** | **54.76/\*\*\*** | 0.09/0.76 |
| Femur Length | **307.27/\*\*\*** | **59.16/\*\*\*** | 2.19/0.14 |
| Tibia Length | **140.64/\*\*\*** | **20.94/\*\*\*** | 0.08/0.76 |

GLM analyses of the relationship between head size and hind limb length reveal largely concordant patterns regardless of which metric (head length or post-orbital width) is used to quantify head size (Table 3; supplemental materials). For both measurements, the top model (lowest AICc score) was the one containing a different intercept of the relationship between head size and limb length for the two species, but without a difference in slope (i.e., no interaction between species identity and the head size/SVL relationship). These top models also include no effect of sex on the relationship between head size and limb length, but in both cases the model that did include sex was also within or nearly within the set of credible models (deltaAIC of 1.52 for head length, and deltaAIC of 2.2 for post-orbital width).

## Discussion

*Hemidactylus mabouia* ranks among the most pervasive invasive lizard species in the neotropics (Rödder et al. 2008; Weterings and Vetter 2018). This species has repeatedly been hypothesized to represent a superior competitor that restricts access to food resources (Rocha et al. 2011; Hughes et al. 2015; Williams et al. 2016) and thereby promotes the extirpation of native, as well as non-native, geckos (van Buurt 2004; van Buurt 2006). Our study provides support for this hypothesis, showing that on Curaçao, *H. mabouia* not only competes with the native gecko *Phyllodactylus martini* for prey resources but preys upon this and other vertebrate species. Notably, both stable isotopic and stomach contents demonstrate that *H. mabouia* will readily consume vertebrate prey items that include *P. martini*, *Gonatodes antillensis* (the Venezuelan Coastal Clawed Gecko), and the non-native blind snake *Ramphotyphlops braminus*. Additionally, we demonstrate larger sizes in feeding associated traits and limb lengths that may be advantageous for *H. mabouia* during the rapid forward propulsive locomotion associated with ambush predation. Given the ubiquity of *H. mabouia* throughout the neotropics, our results provide a new perspective for understanding the complexity of *Hemidactylus* spp. invasions, suggesting their potential impact to be vastly underestimated.

### On the competitive advantages of Hemidactylus

Prior work has suggested that *H. mabouia* directly competes with *Phyllodactylus martini* for food resources (Hughes et al. 2015), suggesting resource competition is a major driver of *P. martini’s* displacement. Our analyses are consistent with the expectations of a competitive exploitation hypothesis, demonstrating substantial overlap of major invertebrate prey categories between *H. mabouia* and *P. martini* when the two species co-occur (Fig. 3; Supplemental materials). These prey categories largely reflect common groups of invertebrates associated with human dwellings and artificial lighting in Curaçao (Dornburg et al. 2016) and are consistent with studies of the diet of *H. mabouia* in other urbanizing areas (Bonfiglio et al. 2006; Iturriaga and Marrero 2013; Drüke and Rödder 2017). In addition to dietary overlap, apotential explanation for the dissimilarity in number of prey items found in *H. mabouia* and *P. martini* may stem from the ambush prey capture tactics of *H. mabouia*. Fragments of presumably larger prey items such as roaches, beetles, and spiders were often found in the stomachs of *H. mabouia* in comparison with *P. martini*. Additionally, high numbers of isopods were found in some individuals. This suggests that *H. mabouia* could be opportunistically feeding on larger prey as well prey encountered in daytime refugia. The latter could also explain the finding of a blind snake within an individual *H. mabouia*. While partially digested fragments of invertebrate body parts prohibit further testing of whether *H. mabouia* is more effectively harnessing larger prey, this hypothesis raises several possibilities of how the natural history of these species influences differential patterns of foraging and prey capture.

There are different responses to bright lighting between these species with *Hemidactylus mabouia* readily foraging directly at brightly lit artificial lights (Perry and Fisher 2006; Hughes et al. 2015). This strategy reduces the energetic cost of finding prey as *H. mabouia* can harness the potential of artificial lights as a lure for attracting large prey resources (Gaston et al. 2013) while simultaneously gaining a potential thermal advantage (Perry et al. 2008). In contrast, *P. martini* avoids direct bright lights, and is often found foraging along the more dimly lit periphery of buildings (Hughes et al. 2015). As such, *P. martini* may have an ecological disadvantage to *H. mabouia*, as the former may need to spend more time locating prey. Furthermore, this small change in prey foraging may put *P. martini* in contact with arthropod vectors for nematodes not encountered by *H. mabouia*, as suggested by our observation of a difference in parasite infestations between species. Given that lizards are often transport hosts for mammalian parasites (Incedogan et al 2014; Dornburg et al 2019), including nematodes (Goldberg and Bursey 2000), further testing of differences in parasite frequencies between *Hemidactylus* and its native competitors represents an exciting direction additional research of high relevance to animal health.

In addition to having an advantage in light tolerance, our analyses of trait morphological variation suggest that *H. mabouia* has a size advantage over *P. martini*, possessing overall larger size, as well as larger heads, hind limbs, and other traits (Fig. 4 & Table 2). Increases in head height and head length are associated with increases in bite force and more efficient prey capture in geckos (Cameron et al. 2013; Massetti et al. 2017), as well as other lizard species (Verwaijen et al. 2002; Dufour et al. 2018). Functionally, this advantage is thought to arise by the combination of increasing space to accommodate increases in mandible adductor muscle sizes as well as changes in attachment angle that provide force advantages (Herrel et al. 2001). Head sizes were larger in both male and female *H. mabouia* relative to *P. martini*, with only head width (temporal width) significantly different between the sexes (Supplemental materials). Males of the closely related *Hemidactylus turcicus* have also been found to have larger head widths that are hypothesized to be the result of sexual selection (Iturriaga and Marrero 2013), and our results suggest a similar pattern of dimorphism occurs in *H. mabouia*. However, larger heads also come at a cost. Increased head sizes can negatively impact sprinting speed in lizards (Cameron et al. 2013), and our additional finding of increased hind limb lengths in both sexes of *H. mabouia* may reflect the species avoiding a fundamental locomotor trade-off (Table 3, Supplemental materials). A similar compensation has been reported in *Hemidactylus frenatus* (Cameron et al. 2013) suggesting this is potentially a general feature of *Hemidactylus* locomotor morphology.

*Hemidactylus mabouia* has been subjected to an unintentional experiment of introduction to human mediated landscapes across the new world for centuries (Goeldi 1902; Van Buurt 2004; Carranza and Arnold 2006). But, whether colonization of human structures has placed this species under selection for changes in locomotor morphology remains unclear. Longer hind limbs in lizards are often correlated with increased sprint speeds and forward propulsion in lizards (Bonine and Garland 1999; Cameron et al. 2013; Winchell et al 2018), thereby providing an advantage for an ambush predator such as *H. mabouia* relying on a combination of ambush and pursuit to capture prey. Additionally, recent work placing front limbs into the context of gecko locomotion models (Birn-Jeffery and Higham 2016; Zhuang and Higham 2016) provides strong evidence that locomotor function is decoupled between fore- and hind limbs. In contrast to hind limbs, which act as primary axes of propulsion, front limbs are primarily used for braking and downward locomotion (Birn-Jeffery and Higham 2016). Quantifications of limb morphology across major lineages of geckos suggest shorter front limbs relative to hindlimbs to be a hallmark of gecko locomotor morphology, with all species having between a 10 to 35% reduction in front limb proportions (Hagey et al. 2017), a finding consistent with our analysis *H. mabouia* limb proportions (Supplemental materials).

Primarily shorter front limbs could shorten the swing time, thereby aiding in maintaining speed and stance in downward movements (Birn-Jeffery and Higham 2016). Our finding of shorter front to hindlimbs in *H. mabouia* are consistent with expectations of selection for locomotion on steeply inclined surfaces such as walls that is coupled with large hindlimbs for sprinting. But, the significant negative scaling relationship between forelimb length and body size for *P. martini* also highlights the potential that additional major differences in locomotor mode and performance between these species exist. Currently, the foraging mode and activity patterns of *P. martini* remain little studied, as do those of *H. mabouia* in their native range. As comparative studies of gecko functional locomotor morphology and performance continue to illuminate the role of forelimbs in gecko locomotor morphology, future comparisons of locomotor morphology and performance between and within these species offers a promising and exciting research frontier.

### The role of predation in Hemidactylus invasions?

Superiority in food resource competition has repeatedly been hypothesized as a major factor facilitating the establishment of *H. mabouia* at the expense of native geckos (Petren and Case 1996; Hoskin 2011; Hughes et al. 2015). Our stomach content analyses revealed a significant overlap of major invertebrate prey resources, thereby supporting expectations of food resource competition (Table 1; Fig. 2 & 3). However, our study additionally provides direct evidence from stomach contents and indirect evidence from the analysis of ∂15N isotopes that *Hemidactylus mabouia* acts as an opportunistic vertebrate predator. These provide a broader context to previously reported single instances of predation by *Hemidactylus mabouia* on *Phyllodactylus martini* (Dornburg et al. 2016) and *Gonatodes antillensis* (Dornburg et al. 2011), as well as observations of cannibalism (Bonfiglio et al. 2006). We additionally report the first instance of ophiophagy in *H. mabouia* (Table 1), suggesting that this species readily consumes smaller vertebrate prey. This raises a question: How frequent are such predation events?

Our isotopic analyses provide some insights to this question, indicating numerous individual *H. mabouia* are feeding at a trophic level higher than *P. martini*. As vertebrates represent the only consumers of a higher trophic position in Curaçao, ∂15N values suggest that predation of vertebrates by *H. mabouia* may not be rare events. Investigations of feeding ecology of the closely related *H. frenatus* have also reported intraspecific juvenile predation (Hunsaker 1966; Bolger and Case 1992; Case et al. 1994), suggesting *H. mabouia* to be similarly opportunistic. Juvenile *H. mabouia* are known to avoid predation by larger adults by foraging low to the ground (Howard et al. 2001). It is likely that our results reflect a signature of juvenile mortality of *P. martini* as juveniles of this species will readily forage across a range of wall elevations including those that are occupied by *H. mabouia* (van Buurt 2004). In addition to lowering recruitment, predation on juvenile *P. martini* could offer a competitive advantage for juvenile *H. mabouia*. By reducing the density of interspecific competitors at the juvenile stage, more juvenile *H. mabouia* would be able to transition to adult and more swiftly increase overall population sizes. As *H. mabouia* readily achieves high carrying capacities that can exceed those of other *Hemidactylus* species (Short and Petren 2011), this in turn could greatly increase the pressure of additional density dependent effects on the persistence of native species.

Evidence for predation of smaller vertebrates by *H. mabouia* raises the concern that in addition to displacing populations of native geckos, the presence of *H. mabouia* can negatively impact overall population structure. Although demographic studies of geckos impacted by *H. mabouia* have been limited, analyses of *Phyllodactylus tuberculosis* in Mexico have implicated the presence of *H. mabouia* in severe contractions of effective population size and recent genetic bottlenecks (Blair et al. 2015). We argue that further assessing the role of *H. mabouia* in juvenile survivorship represents an important, but currently neglected aspect of this species invasion biology. These studies are of particular importance as *H. mabouia* is increasingly being found in non-urban areas throughout its invaded range (Rocha et al. 2011), challenging the assumption that this invasion is limited to urbanizing areas. Fortunately for Curaçao and other similar desert habitats, invasion into the native bush habitat may not be possible due to mechanical properties of *H. mabouia* toe pad adhesion. *Hemidactylus* geckos all possess a basal toe pad system that may not be capable of successfully gaining traction on the loose, and dusty, rocky soil of the island (Russell and Delaugerre 2017). This hypothesis remains to be tested, but if it is supported, the impact of predation and competition on native gecko population sizes in dusty, arid environments could be mitigated by integrating continual corridors of native habitat into urban planning efforts. Such efforts would yield ‘enemy-free’ space and thereby increase the probability of the long-term persistence of native gecko species (Cole et al. 2005).

## Acknowledgements

We thank Mark Vermeij and the entire staff of the Caribbean Research and Management of Biodiversity foundation (CARMABI) for facilitating collection efforts and export permitting, Sunshine and David at Curaçao Sunshine Getaways for enabling us to convert the second floor of their home into a field station, and the Westpunt Syndicate for their continual field assistance. Specimens were collected under the approved guidelines of IACUC (Institutional Animal Care and Use Committee) protocol 2012-10681 (Ichthyology and Herpetology at the Yale Peabody Museum of Natural History). All data and analysis scripts associated with this study have been archived on Zenodo (doi pending acceptance). We also would like to thank E. Ferraro, Enie Hensel, and two anonymous referees for excellent feedback on previous versions of this manuscript.

## Literature Cited

Allen M, Poggiali D, Whitaker K, Marshall TR, Kievit, RA (2019) Raincloud plots: a multi-platform tool for robust data visualization. Wellcome open research 4:63

González-Oreja JA, Zuria I, Carbo-Ramirez P, Charre GM (2018) Using variation partitioning techniques to quantify the effects of invasive alien species on native urban bird assemblages. Biol Invasions 20:2861–2874

Bateman PW, Fleming PA (2012) Big city life: carnivores in urban environments. J Zool 287:1–23

Birn-Jeffery AV, Higham TE (2016) Geckos decouple fore- and hind limb kinematics in response to changes in incline. Front Zool 13:11

Blair C, Jiménez Arcos VH, de la Cruz FRM, Murphy RW (2015) Historical and contemporary demography of leaf-toed geckos (Phyllodactylidae: *Phyllodactylus tuberculosus saxatilis*) in the Mexican dry forest. Conservation Genetics 16:419–429

Bolger DT, Case TJ (1992) Intra- and interspecific interference behaviour among sexual and asexual geckos. Animal Behaviour 44:21–30

Bonfiglio F, Balestrin RL, Cappellari LH (2006) Diet of *Hemidactylus mabouia* (Sauria, Gekkonidae) in urban area of southern Brazil. Biociências 14:107–111

Bonine KE, Garland Jr T (1999) Sprint performance of phrynosomatid lizards, measured on a high-speed treadmill, correlates with hindlimb length. Journal of Zoology, 248:255–265

Burgos-Rodríguez JA, Avilés-Rodríguez KJ, Kolbe JJ (2016) Effects of invasive Green Iguanas *(Iguana iguana*) on seed germination and seed dispersal potential in southeastern Puerto Rico. Biological Invasions 18:2775–2782

Burnham KP, Anderson DR (2004) Multimodel inference: understanding AIC and BIC in model selection. Sociological methods & research, 33:261–304

Buzan E (2017) Changes in rodent communities as consequence of urbanization and inappropriate waste management. Appl Ecol Environ Res 15:573–588

Cameron SF, Wynn ML, Wilson RS (2013) Sex-specific trade-offs and compensatory mechanisms: bite force and sprint speed pose conflicting demands on the design of geckos (*Hemidactylus frenatus*). J Exp Biol 216:3781–3789

Capinha C, Seebens H, Cassey P, García-Díaz P, Lenzner B, Mang T, Moser D, Pyšek P, Rödder D, Scalera R, Winter M, Dullinger S, Essl F (2017) Diversity, biogeography and the global flows of alien amphibians and reptiles. Diversity and Distributions 23:1313–1322

Carranza S, Arnold EN (2006) Systematics, biogeography, and evolution of Hemidactylus geckos (Reptilia: Gekkonidae) elucidated using mitochondrial DNA sequences. Mol Phylogenet Evol 38:531–545

Case TJ, Bolger DT, Petren K (1994) Invasions and Competitive Displacement among House Geckos in the Tropical Pacific. Ecology 75:464–477

Chapman MG, Underwood AJ (1999) Ecological patterns in multivariate assemblages:information and interpretation of negative values in ANOSIM tests. Mar Ecol Prog Ser 180:257–265

Clarke KR (1993) Non-parametric multivariate analyses of changes in community structure. Aust J Ecol 18:117–143

Cole NC, Jones CG, Harris S (2005) The need for enemy-free space: The impact of an invasive gecko on island endemics. Biological Conservation 125:467–474

Donald GL, Paterson CG (1977) Effect of preservation on wet weight biomass of chironomid larvae. Hydrobiologia 53:75–80

Donihue CM, Herrel A, Fabre A-C, Kamath A, Geneva AJ, Schoener TW, Kolba JJ, Losos JB (2018) Hurricane-induced selection on the morphology of an island lizard. Nature 560:88–91

Dornburg A, Lippi C, Federman S, et al (2016) Disentangling the Influence of Urbanization and Invasion on Endemic Geckos in Tropical Biodiversity Hot Spots: A Case Study of *Phyllodactylus martini* (Squamata: Phyllodactylidae) along an Urban Gradient in Curaçao. Bulletin of the Peabody Museum of Natural History 57:147–164

Dornburg A, Warren DL, Iglesias T, Brandley MC (2011) Natural History Observations of the Ichthyological and Herpetological Fauna on the Island of Curaçao (Netherlands). Bulletin of the Peabody Museum of Natural History 52:181–186

Dornburg A, Lamb AD, Warren D, Watkins-Colwell GJ, Lewbart GA, Flowers J (2019). Are Geckos Paratenic Hosts for Caribbean Island Acanthocephalans? Evidence from *Gonatodes antillensis* and a Global Review of Squamate Reptiles Acting as Transport Hosts. Bulletin of the Peabody Museum of Natural History, 60:55–79.

Drüke Y, Rödder D (2017) Feeding ecology of the invasive gecko species *Hemidactylus mabouia* (Moreau de Jonnès, 1818) (Sauria: Gekkonidae) in São Sebastião (Brazil). Bonn Zool Bull 66:85–93

Dufour CMS, Losos JB, Herrel A (2018) Do differences in bite force and head morphology between a native and an introduced species of anole influence the outcome of species interactions? Biological Journal of the Linnean Society XX: 1–10

Falcón W, Ackerman JD, Recart W, Daehler CC (2013) Biology and Impacts of Pacific Island Invasive Species. 10. *Iguana iguana*, the Green Iguana (Squamata: Iguanidae). Pac Sci 67:157–186

Gaston KJ, Bennie J, Davies TW, Hopkins J (2013) The ecological impacts of nighttime light pollution: a mechanistic appraisal. Biol Rev Camb Philos Soc 88:912–927

Goeldi EA (1902) Lagartos do Brazil. Boletim do Museu Paraense 3:499–560

Goldberg SR, Bursey CR (2000) Transport of helminths to Hawaii via the brown anole, *Anolis sagrei* (Polychrotidae). Journal of Parasitology 86:750–756.

Hagey TJ, Harte S, Vickers M, et al (2017) There’s more than one way to climb a tree: Limb length and microhabitat use in lizards with toe pads. PLoS One 12:e0184641

Herrel A, De Grauw E, Lemos-Espinal JA (2001) Head shape and bite performance in xenosaurid lizards. J Exp Zool 290:101–107

Hintze JL, Nelson RD (1998) Violin Plots: A Box Plot-Density Trace Synergism. The American Statistician 52:181

Hoskin CJ (2011) The invasion and potential impact of the Asian House Gecko (*Hemidactylus frenatus*) in Australia. Austral Ecology 36:240–251

Howard KG, Parmerlee JS, Powell R (2001) Natural history of the edificarian geckos *Hemidactylus mabouia*, *Thecadactylus rapicauda*, and *Sphaerodactylus sputator* on Anguilla. Caribb J Sci 37:285–287

Hughes DF, Meshaka WE Jr, van Buurt G (2015) The superior colonizing gecko *Hemidactylus mabouia* on Curaçao: conservation implications for the native gecko *Phyllodactylus martini*. J Herpetol 49:60–63

Hunsaker D (1966) Notes on the population expansion of the house gecko, *Hemidactylus frenatus*. Philipp J Sci 95:121–122

Incedogan S, Yildirimhan HS, Bursey CR, 2014. Helminth parasites of the ocellated skink, *Chalcides ocellatus* (Forskal, 1775) (Scincidae) from Turkey. Comparative Parasitology, 81:260–269

Iturriaga M, Marrero R (2013) Feeding ecology of the Tropical House Gecko *Hemidactylus mabouia* (Sauria: Gekkonidae) during the dry season in Havana, Cuba. Herpetol Notes 6:11–17

Kratochvíl L, Frynta D (2002) Body size, male combat and the evolution of sexual dimorphism in eublepharid geckos (Squamata: Eublepharidae). Biological Journal of the Linnean Society 76:303–314.

Kraus F (2015) Impacts from Invasive Reptiles and Amphibians. Annu Rev Ecol Evol Syst 46:75–97

Lapiedra O, Chejanovski Z, Kolbe JJ (2017) Urbanization and biological invasion shape animal personalities. Glob Chang Biol 23:592–603

Liu C, He D, Chen Y, Olden JD (2017) Species invasions threaten the antiquity of China’s freshwater fish fauna. Diversity and Distributions 23:556–566

Marshall JD, Brooks JR, Lajtha K (2007) Sources of variation in the stable isotopic composition of plants. Stable isotopes in ecology and environmental science 2:22–60

Massetti F, Gomes V, Perera A, et al (2017) Morphological and functional implications of sexual size dimorphism in the Moorish gecko, *Tarentola mauritanica*. Biological Journal of the Linnean Society 122:197–209

McKinney ML (2006) Urbanization as a major cause of biotic homogenization. Biological Conservation 127:247–260

McKinney ML, Lockwood JL (1999) Biotic homogenization: a few winners replacing many losers in the next mass extinction. Trends Ecol Evol 14:450–453

Mooney HA, Cleland EE (2001) The evolutionary impact of invasive species. Proc Natl Acad Sci USA 98:5446–5451

Morais P, Reichard M (2018) Cryptic invasions: A review. Sci Total Environ 613-614:1438–1448

Paini DR, Sheppard AW, Cook DC, et al (2016) Global threat to agriculture from invasive species. Proc Natl Acad Sci USA 113:7575–7579

Pedersen T, Fuhrmann MM, Lindstrøm U, et al (2018) Effects of the invasive red king crab on food web structure and ecosystem properties in an Atlantic fjord. Mar Ecol Prog Ser 596:13–31

Perry G (1996) The evolution of sexual dimorphism in the lizard *Anolis polylepis* (Iguania): evidence from intraspecific variation in foraging behavior and diet. Can J Zool 74:1238–1245

Perry G, Buchanan BW, Fisher RN, et al (2008) Effects of artificial night lighting on amphibians and reptiles in urban environments. Urban herpetology 3:239–256

Perry G, Fisher RN (2006) Night lights and reptiles: observed and potential effects. In: Ecological consequences of artificial night lighting. 169–191

Petren K, Case TJ (1998) Habitat structure determines competition intensity and invasion success in gecko lizards. Proc Natl Acad Sci U S A 95:11739–11744

Petren K, Case TJ (1996) An Experimental Demonstration of Exploitation Competition in an Ongoing Invasion. Ecology 77:118–132

Post DM (2002) Using stable isotopes to estimate trophic position: models, methods, and assumptions. Ecology 83:703–718

R Core Team (2018) R: A language and environment for statistical computing. R Foundation for Statistical Computing, Vienna, Austria

Richmond JQ, Wood DA, Stanford JW, Fisher RN (2015) Testing for multiple invasion routes and source populations for the invasive brown treesnake (*Boiga irregularis*) on Guam: implications for pest management. Biological Invasions 17:337–349

Rocha CFD, Anjos LA, Bergallo HG (2011) Conquering Brazil: the invasion by the exotic gekkonid lizard *Hemidactylus mabouia* (Squamata) in Brazilian natural environments. Zoologia (Curitiba) 28:747–754

Roches SD, Des Roches S, Harmon LJ, Rosenblum EB (2016) Colonization of a novel depauperate habitat leads to trophic niche shifts in three desert lizard species. Oikos 125:343–353

Rodda GH, Savidge JA (2007) Biology and Impacts of Pacific Island Invasive Species. 2. *Boiga irregularis,* the Brown Tree Snake (Reptilia: Colubridae) 1. Pac Sci 61:307–325

Rödder D, Solé M, Böhme W (2008) Predicting the potential distributions of two alien invasive Housegeckos (Gekkonidae: *Hemidactylus frenatus, Hemidactylus mabouia*). North-Western Journal of Zoology 4:236–246

Russell AP, Delaugerre MJ (2017) Left in the dust: differential effectiveness of the two alternative adhesive pad configurations in geckos (Reptilia: Gekkota). Journal of Zoology 301:61–68

Scharf I, Meiri S, (2013) Sexual dimorphism of heads and abdomens: different approaches to ‘being large’ in female and male lizards. Biological Journal of the Linnean Society, 110:665–673.

Shechonge A, Ngatunga BP, Bradbeer SJ, et al (2019) Widespread colonisation of Tanzanian catchments by introduced *Oreochromis* tilapia fishes: the legacy from decades of deliberate introduction. Hydrobiologia 832:235–253

Short KH, Petren K (2011) Rapid species displacement during the invasion of Florida by the tropical house gecko *Hemidactylus mabouia*. Biol Invasions 14:1177–1186

Smith BJ, Cherkiss MS, Hart KM, et al (2016) Betrayal: radio-tagged Burmese pythons reveal locations of conspecifics in Everglades National Park. Biological Invasions 18:3239–3250

Toussaint A, Beauchard O, Oberdorff T, et al (2016) Worldwide freshwater fish homogenization is driven by a few widespread non-native species. Biological Invasions 18:1295–1304

Trentanovi G, von der Lippe M, Sitzia T, et al (2013) Biotic homogenization at the community scale: disentangling the roles of urbanization and plant invasion. Diversity and Distributions 19:738748

Useni Sikuzani Y, Sambiéni Kouagou R, Maréchal J, et al (2018) Changes in the Spatial Pattern and Ecological Functionalities of Green Spaces in Lubumbashi (the Democratic Republic of Congo) in Relation With the Degree of Urbanization. Tropical Conservation Science 11:1940082918771325

van Buurt G (2004) Field Guide to the Reptiles and Amphibians of Aruba, Curacao and Bonaire. Serpents Tale

van Buurt G (2006) Conservation of amphibians and reptiles in Aruba, Curaçao and Bonaire. Applied Herpetology 3:307–321

Verwaijen D, Van Damme R, Herrel A (2002) Relationships between head size, bite force, prey handling efficiency and diet in two sympatric lacertid lizards. Functional Ecology 16:842–850

Vidal MA, Sabat P (2010) Stable isotopes document mainland–island divergence in resource use without concomitant physiological changes in the lizard *Liolaemus pictus*. Comparative Biochemistry and Physiology Part B: Biochemistry and Molecular Biology 156:61–67

Vitt LJ (1983) Tail loss in lizards: the significance of foraging and predator escape modes. Herpetologica 39:151–162

Weber MJ, Brown ML (2009) Effects of Common Carp on Aquatic Ecosystems 80 Years after “Carp as a Dominant” Ecological Insights for Fisheries Management. Rev Fish Sci 17:524–537

Weterings R, Vetter KC (2018) Invasive house geckos (*Hemidactylus* spp.): their current, potential and future distribution. Curr Zool 64:559–573

Wiles GJ, Bart J, Beck RE, Aguon CF (2003) Impacts of the Brown Tree Snake: Patterns of Decline and Species Persistence in Guam’s Avifauna. Conservation Biology 17:1350–1360

Williams R, Pernetta AP, Horrocks JA (2016) Outcompeted by an invader? Interference and exploitative competition between tropical house gecko (*Hemidactylus mabouia*) and Barbados leaf-toed gecko (*Phyllodactylus pulcher*) for diurnal refuges in anthropogenic coastal habitats. Integr Zool 11:229–238

Willson JD (2017) Indirect effects of invasive Burmese pythons on ecosystems in southern Florida. J Appl Ecol 54:1251–1258

Winchell KM, Maayan I, Fredette JR, Revell LJ (2018) Linking locomotor performance to morphological shifts in urban lizards. Proc Biol Sci 285.: doi: 10.1098/rspb.2018.0229

Winchell KM, Reynolds RG, Prado-Irwin SR, et al (2016) Phenotypic shifts in urban areas in the tropical lizard *Anolis cristatellus*. Evolution 70:1009–1022

Yam RSW, Fan Y-T, Wang T-T (2016) Importance of Macrophyte Quality in Determining Life-History Traits of the Apple Snails *Pomacea canaliculata:* Implications for Bottom-Up Management of an Invasive Herbivorous Pest in Constructed Wetlands. Int J Environ Res Public Health 13: doi: 10.3390/ijerph13030248

Young HS, Parker IM, Gilbert GS, et al (2017) Introduced Species, Disease Ecology, and Biodiversity-Disease Relationships. Trends Ecol Evol 32:41–54

Zaaf A, Van Damme R (2001) Limb proportions in climbing and ground-dwelling geckos (Lepidosauria, Gekkonidae): a phylogenetically informed analysis. Zoomorphology 121:45–53

Zhuang MV, Higham TE (2016) Arboreal Day Geckos (*Phelsuma madagascariensis*) Differentially Modulate Fore- and Hind Limb Kinematics in Response to Changes in Habitat Structure. PLoS One 11:e0153520

