## Supplemental Tables and Figures for "What makes *Hemidactylus* invasions successful? A case study on the island of Curaçao"

**Supplemental Table 1 | Museum catalog numbers and collection locality of *Phyllodactylus martini* and *Hemidactylus mabouia* specimens used.** Abbreviations: CARMABI, Caribbean Research and Management of Biodiversity; YPM HERR, Division of Vertebrate Zoology Herpetology Collection (Reptiles), Peabody Museum of Natural History, Yale University.

| **Locality- Habitat Type** | ***Phyllodactylus martini*** | ***Hemidactylus mabouia*** |
|---|---|---|
| *Willemstadt*- Urban | --- | **YPM HERR** 18122, 18124, 18356, 18357, 18358, 18359, 18350, 18351, 18352, 18353, 18354, 18355, 18121, 18122, 18124 |
| *Saint Anna Bay* - Urban | --- | **YPM HERR** 18151, 18152, 18154 |
| *CARMABI*- Edge | **YPM HERR** 18437, 18435, 18436 | **YPM HERR** 18121, 18376, 18434 |
| *Lagun*- Edge | **YPM HERR** 18462, 18463, 18466, 18461, 18467, 18626 | **YPM HERR** 18464, 18470, 18478, 18475, 18474, 18472, 18471, 18465, 18468, 18473, 18476, 18469 |
| *Westpunt*- Edge | **YPM HERR** 17585, 17586, 17587, 17588, 17589, 17590, 17591, 17592, 17593, 17594, 17595, 17596, 17597, 17598, 17600, 17601, 17602, 17604, 17607, 18168, 18169, 18170, 18171, 18172, 18173, 18175, 18176, 18177, 18179, 18180, 18141, 18183, 18184, 18165, 18167, 18185, 18186, 18187, 18616, 18612, 18617, 18614, 18645, 18646, 18622, 18623, 18611, 18619 | **YPM HERR** 18133, 18134, 18135, 18136, 18137, 18140, 18127, 18142, 18143, 18144, 18632, 18630, 18633, 18631, 18638, 17557, 17563, 17564, 17565, 17566, 17567, 17568, 17569, 17570, 17571, 17572, 17573, 17575, 17576, 17577, 17578, 18127, 18127, 18129, 18130, 18131, 18132, 18133, 18134, 18135, 18136, 18137, 18138, 18139, 18140, 18142, 18143, 18144, 18145, 18146, 18149, 18150, 18151, 18152, 18154, 18155, 18631 |
| *Shete Boca*- Rural | **YPM HERR** 18342, 18343, 18344, 18345, 18345, 18347, 18348, 18349, 18629, 18625, 18624, 18627, 18628, 18626 | --- |

**Supplemental Table 2 | ANCOVA results testing the effect of snout-vent length (SVL), species, and their interaction on measured morphological characters with non-significant interactions removed.** Bolded values indicate significant effects. * stands for *P*-values ranging from 0.05 to 0.01, ** for *P*-values ranging from 0.01 to 0.001 and *** for *P*-values smaller than 0.001.

| Trait | log(SVL) (F/P) | Species (F/P) | Species:log(SL) (F/P) |
|---|---|---|---|
| Post Orbit Width | **122.67/*** ** | **6.02/* ** | --- |
| Temporalis Width | **1159.48/*** ** | **32.68/*** ** | --- |
| Head Length | **1307.8/*** ** | **102.6/*** ** | --- |
| Jaw Length | **530.02/*** ** | **86.69/*** ** | --- |
| Head Height | **359.84/*** ** | 3.248/0.0738 | --- |
| Humerus Length | **163.53/*** ** | **43.521/*** ** | **8.302/** ** |
| Radius Length | **468.16/*** ** | **55.14/*** ** | --- |
| Femur Length | **304.54 /*** ** | **58.63/*** ** | --- |
| Tibia Length | **141.61/*** ** | **21.09/*** ** | --- |

**Supplemental Table 3 | ANCOVA coefficients from tests of the effect of snout-vent length (SVL), species, and their interaction on measured morphological characters.**

| Trait | Intercept | log(SVL) | Species | Species:log(SVL) |
|---|---|---|---|---|
| Post Orbit Width | -1.666 | 0.841 | -0.070 | 0.050 |
| Temporalis Width | -1.411 | 0.945 | -0.053 | 0.018 |
| Head Length | -0.440 | 0.820 | -0.074 | 0.015 |
| Jaw Length | -0.683 | 0.744 | -0.177 | 0.033 |
| Head Height | -2.405 | 1.043 | 0.341 | -0.076 |
| Humerus Length | -2.607 | 1.039 | 1.826 | -0.488 |
| Radius Length | -1.752 | 0.935 | -0.160 | 0.0306 |
| Femur Length | -1.467 | 0.927 | 0.624 | -0.175 |
| Tibia Length | -1.692 | 0.879 | 0.150 | -0.050 |

**Supplemental Table 4 | ANCOVA coefficients from tests of the effect of snout-vent length (SVL), species, and their interaction on measured morphological characters with non-significant interactions removed.**

| Trait | Intercept | log(SVL) | Species | Species:log(SVL) |
|---|---|---|---|---|
| Post Orbit Width | -1.713 | 0.853 | 0.128 | --- |
| Temporalis Width | -1.428 | 0.949 | 0.016 | --- |
| Head Length | -0.454 | 0.823 | -0.167 | --- |
| Jaw Length | -0.714 | 0.752 | -0.047 | --- |
| Head Height | -2.333 | 1.025 | 0.043 | --- |
| Humerus Length | -2.607 | 1.039 | 1.826 | -0.488 |
| Radius Length | -1.781 | 0.941 | -0.040 | --- |
| Femur Length | -1.300 | 0.885 | -0.065 | -0.175 |
| Tibia Length | -1.644 | 0.867 | -0.048 | -0.050 |

**Supplemental Table 5 | Principal component factor loadings by axis for morphological traits of *Phyllodactylus martini* and *Hemidactylus mabouia*.** Top contributing traits indicated in bold for each axis. Abbreviations: PrC = Proportional contribution to PC axis.

| Trait | PC1 \| $(PC1)^2$ \| PrC | PC2 \| $(PC2)^2$ \| PrC | PC3 \| $(PC3)^2$ \| PrC |
|---|---|---|---|
| Post orbital width | **0.581 \| 0.337 \| 0.235** | -0.307 \| 0.094 \| 0.125 | 0.195 \| 0.038 \| 0.081 |
| Temporalis width | 0.177 \| 0.031 \| 0.072 | -0.318 \| 0.101 \| 0.130 | 0.304 \| 0.092 \| 0.127 |
| Head length | 0.083 \| 0.007 \| 0.034 | **-0.700 \| 0.487 \| 0.285** | -0.266 \| 0.071 \| 0.111 |
| Head height | -0.071 \| 0.005 \| 0.029 | -0.094 \| 0.009 \| 0.038 | **0.790 \| 0.623 \| 0.329** |
| Jaw length | -0.145 \| 0.021 \| 0.059 | -0.407 \| 0.166 \| 0.166 | -0.209 \| 0.044 \| 0.087 |
| Humerus length | -0.142 \| 0.020 \| 0.058 | -0.218 \| 0.047 \| 0.089 | -0.044 \| 0.002 \| 0.018 |
| Radius length | -0.302 \| 0.091 \| 0.122 | -0.108 \| 0.012 \| 0.044 | -0.224 \| 0.050 \| 0.093 |
| Femur length | **-0.581 \| 0.337 \| 0.235** | -0.289 \| 0.084 \| 0.118 | 0.261 \| 0.068 \| 0.109 |
| Tibia length | -0.386 \| 0.150 \| 0.157 | 0.012 \| 0.01e-3 \| 0.005 | 0.107 \| 0.012 \| 0.045 |
| *Percent variance explained* | 34.8% | 16.9% | 12.4% |

**Supplemental Figure Captions**

**Supplemental Figure 1 | Results from a resampling procedure assessing differences in morphospace occupation between *Phyllodactylus martini* and *Hemidactylus mabouia* under different levels of individual sampling.** Y axis represents the ratio of morphospace area (*A*) between *H. mabouia and P. martini*. X axis indicates the number of individuals drawn from our dataset at random for (**A)** PC1 & PC2; **(B)** PC2 & PC3; and **(C)** PC1 & PC3. Gray area represents the 25 and 75% quantiles of the resampled morphospace differences, interpolated between sample size replicates. Yellow dots represent the mean of the area ratio for each sample size replicate, with a spline interpolation (yellow line) between points. Dotted line indicates the area ratio quantified when all samples are included in the morphospace (Main Text Figure 4).

**Supplemental Figure 2 | Visualization of stomach content data by species and habitat type.** Rural habitat represents the uninhabited native mondi habitat of the island and only *Phyllodactylus martini* (n=14) were sampled. Urban habitat represents the densely populated areas of the island characterized primarily by the abundance of human-build structures (buildings, roads, etc) and only *Hemidactylus mabouia* (n=14) were sampled. Edge habitat represents areas of the island that are at the interface between, suburbs and the native mondi. For these sites both *P. martini* (n=58) and *H. mabouia* (n=45) were sampled. Columns in sphere correspond to the relative frequency of an individual's prey items. Colors correspond to matching prey categories in legend.

**Supplemental Figure 3A | Feeding-associated trait comparison between *Phyllodactylus martini* (yellow) and *Hemidactylus mabouia* (brown).** Differences in morphological trait size between sexes and species (top) and results from simple linear regression (SLR) between snout to vent length (SVL) and morphological traits (bottom). Traits (from right to left, top to bottom): temporalis width, head length, jaw length, head height, post-orbital width. All lengths in log(mm). Box plot shadings correspond to species + male/female designations in the figure.

**Supplemental Figure 3B | Locomotion-associated trait comparison between *Phyllodactylus martini* (yellow) and *Hemidactylus mabouia* (brown).** Differences in locomotion-associated morphological trait size between sexes and species (top) and results from simple linear regression (SLR) between snout to vent length (SVL) and morphological traits (bottom). All lengths in log(mm). Traits include (from left to right, top to bottom) humerus length, radius length, femur length, tibia length, total upper limb length (humerus+radius), and total lower limb length (femur+tibia). Box plot shadings correspond to species + male/female designations in the figure. Gray shading in the temporalis width plot indicates a statistically significant difference in male and female temporalis width.

#### Supplementary Figure 1

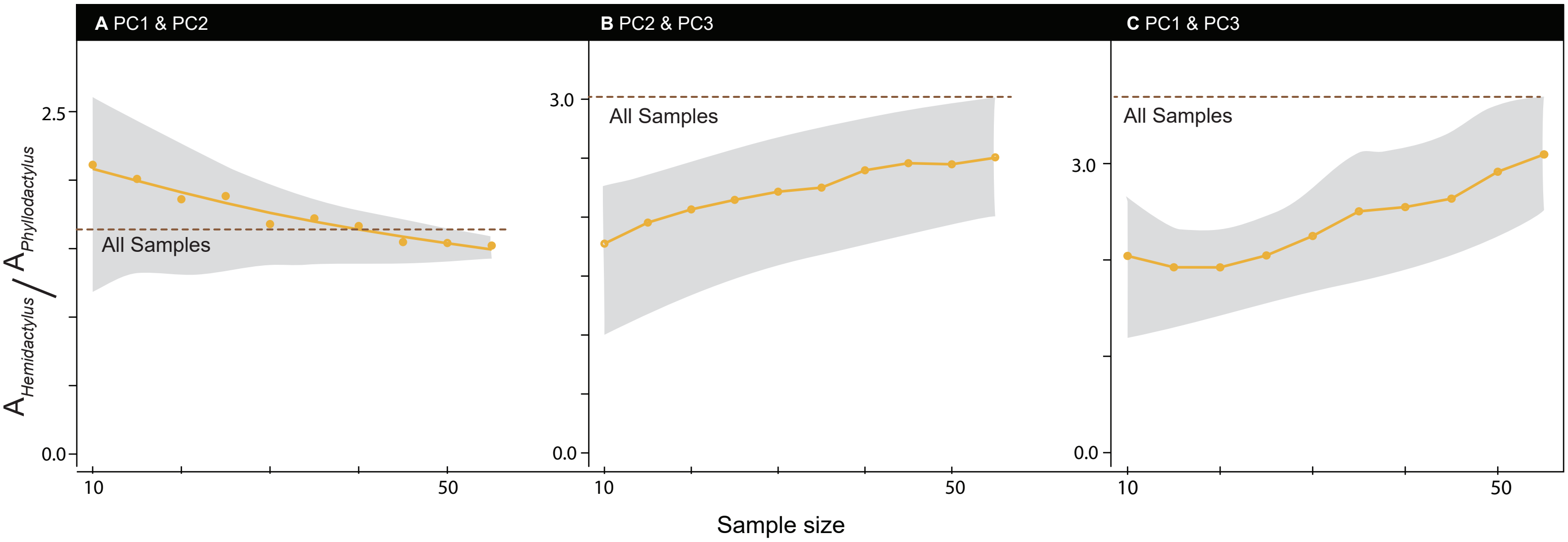

### Supplementary Figure 2

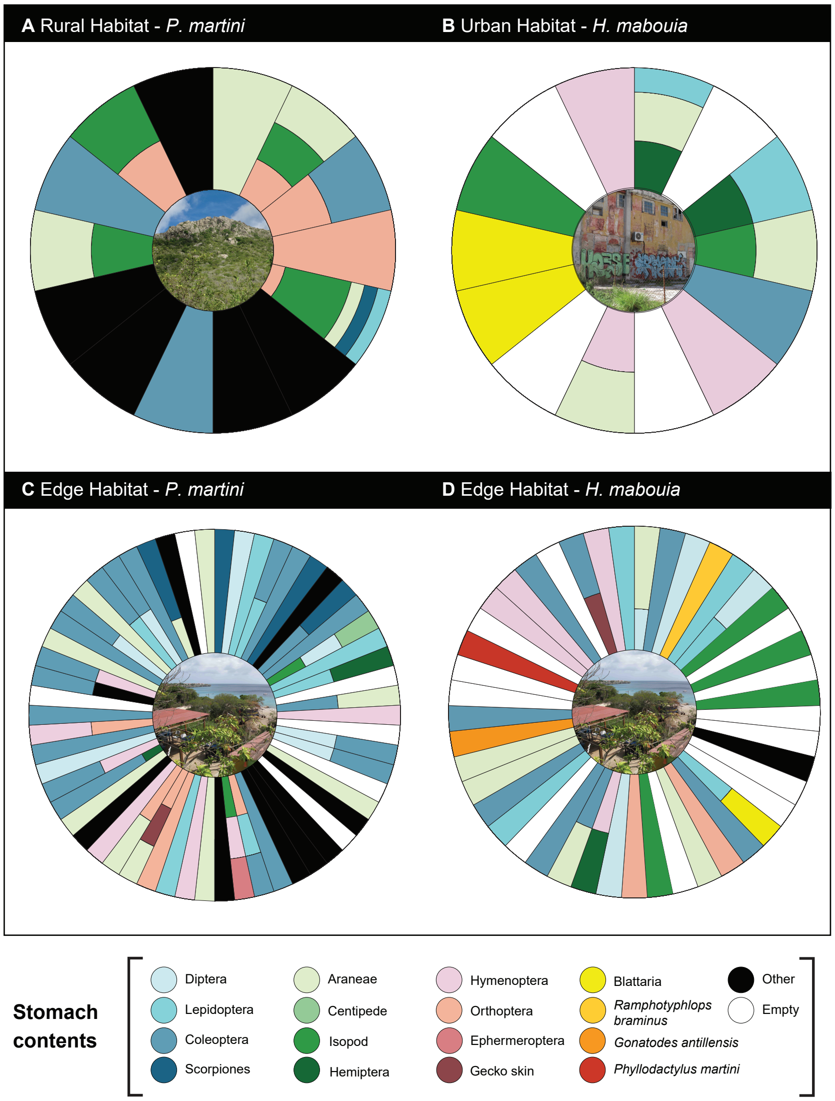

### Supplementary Figure 3A

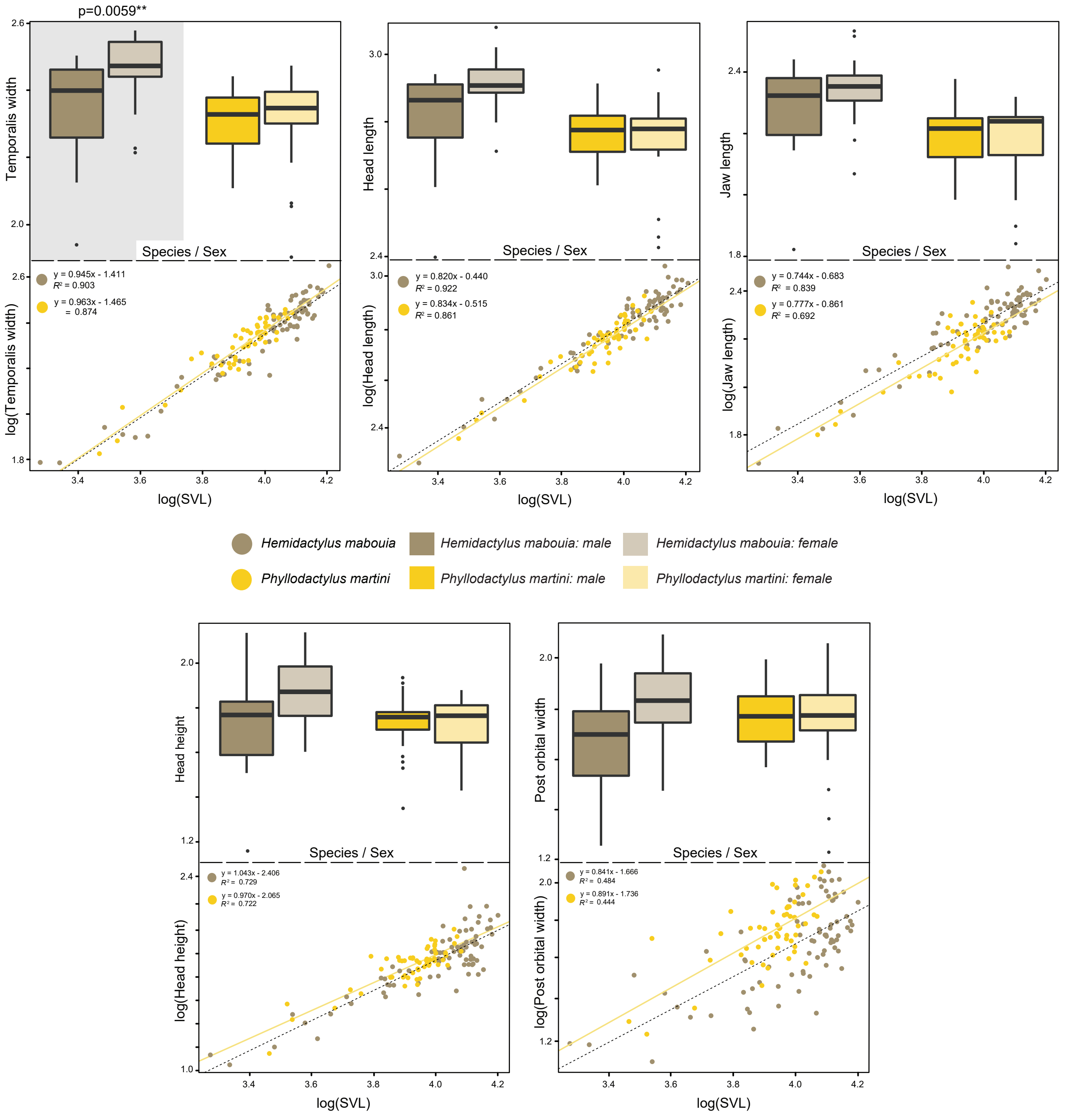

### Supplementary Figure 3B

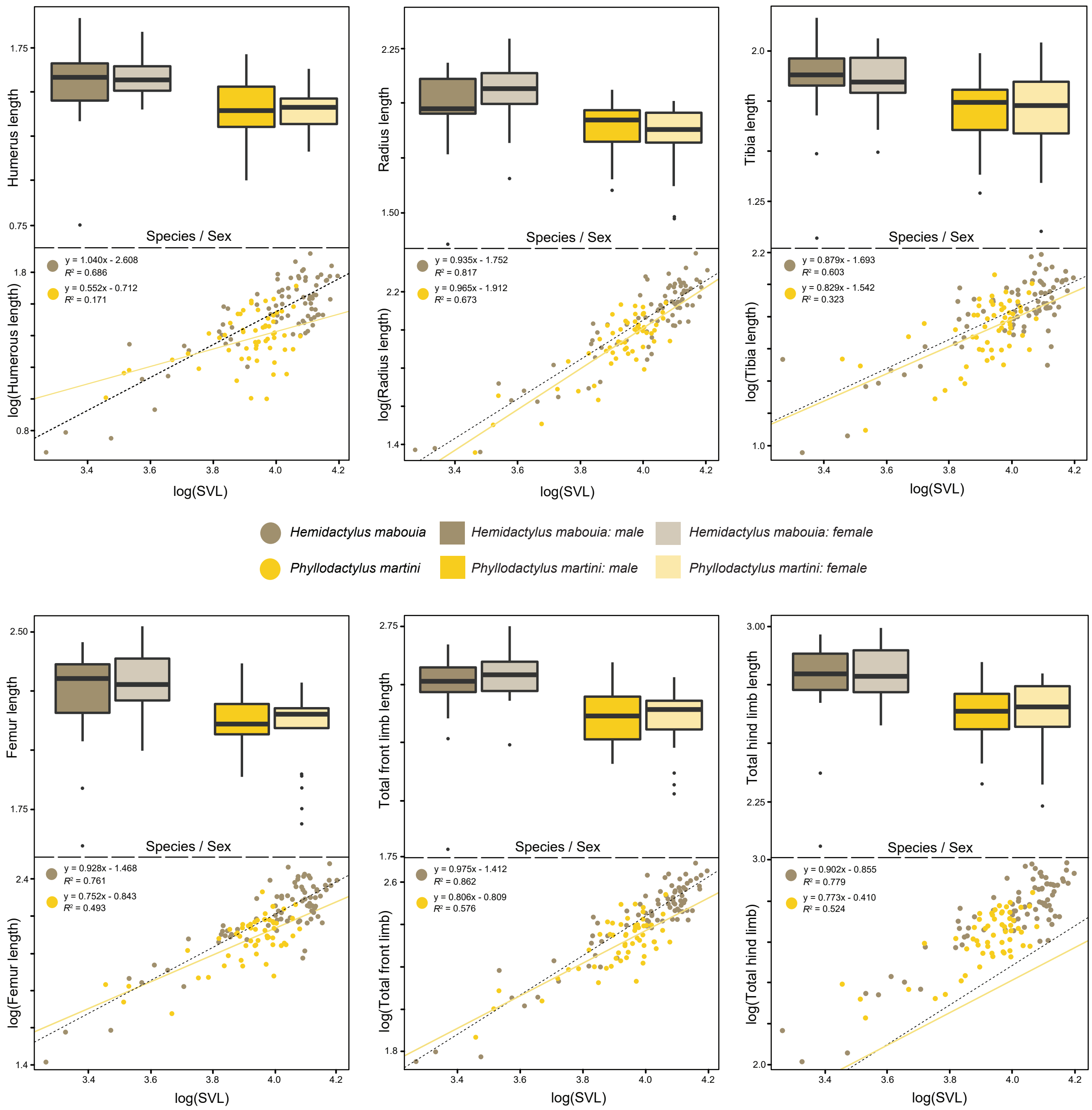
